# Vimentin Intermediate Filaments and Filamentous Actin Form Unexpected Interpenetrating Networks That Redefine the Cell Cortex

**DOI:** 10.1101/2021.07.29.454155

**Authors:** Huayin Wu, Yinan Shen, Suganya Sivagurunathan, Miriam Sarah Weber, Stephen A. Adam, Jennifer H. Shin, Jeffrey J. Fredberg, Ohad Medalia, Robert Goldman, David A. Weitz

## Abstract

Vimentin intermediate filaments (VIFs) and F-actin are filamentous cytoskeletal proteins generally thought to form completely independent networks that have vastly different properties and functions. Here, we show that, unexpectedly, there exist both extensive structural and functional interactions between VIFs and F-actin. We show that VIFs and F-actin form an interpenetrating network (IPN) within the cell cortex and interact synergistically at multiple length scales. This IPN structure has important functional consequences in cells: The IPN results in enhanced contractile forces in the cell. In addition, VIFs influence the diffusive behavior of actin monomers, suggesting specific associations between actin and vimentin proteins in the cytoplasm; this facilitates formation of the IPN and has downstream effects on other actin-driven processes. The results suggest that contributions of VIFs and F-actin are strongly correlated. Such interactions counter generally accepted behavior and are broadly significant given the wide range of processes currently attributed to F-actin alone.

## Introduction

The cytoskeleton is a highly dynamic structure comprised of multiple types of filamentous proteins. In eukaryotic cells, actin, microtubules, and intermediate filaments (IFs) each form intricate networks of entangled and crosslinked filaments. The organization of each individual network is precisely controlled to enable essential cellular functions. However, many core processes also require interactions among the different cytoskeletal components. For example, filamentous-actin (F-actin) and microtubules work together to control cell shape and polarity, which are critical for development, cell migration, and division. Close associations between microtubules and vimentin intermediate filaments (VIFs) have also been proposed based on similarities in their spatial distributions and the dependence of the organization of VIF networks on the microtubule-associated motors, kinesin and dynein (Helfand et al., 2002; Leduc and Etienne-Manneville, 2017; Prahlad et al., 1998). Indeed, there is some experimental evidence that microtubules can template VIF assembly and that VIFs can guide microtubules (Gan et al., 2016; Shabbir et al., 2014). Despite such interactions, it is generally thought that F-actin and VIFs form two co-existing but separate networks. For example, fluorescence microscopy typically reveals the strongest signals for F-actin in the cell periphery, whereas the strongest signals for VIFs are near the nucleus in the bulk cytoplasm, suggesting that the two networks have little or no interaction. Furthermore, the functions of F-actin and VIFs appear to be largely contrasting: F-actin generates forces, whereas VIFs provide stability against these forces. Nevertheless, some evidence suggests there may be connections between vimentin and actin: for example, vimentin knockout cells are less motile and less contractile than their wild-type counterparts (Eckes et al., 1998). Furthermore, some interactions have been observed between actin filaments and the C terminus of vimentin (Duarte et al., 2019, Wirtz 2006) as well as the precursors to keratin, another IF system (Duarte et al., 2019; Esue et al., 2006; Kolsch et al., 2009). These findings suggest that direct interactions or connections may exist between VIFs and F-actin. Such connections would belie our current understanding of the two independent cytoskeletal networks but could have a profound effect on the mechanical properties of cells. The possibility of such connections demands a closer investigation of both the structural and functional interplay between the F-actin and VIF cytoskeletal networks.

Here we present evidence that VIFs and F-actin do work synergistically and form an interpenetrating network (IPN) structure within the cell cortex, defined as the cortical cytoplasm adjacent to the cell surface. In mouse embryonic fibroblasts (MEFs), we observe coupling between F-actin and VIF structures within the cortex, contrary to the widely accepted view that they are each spatially segregated. In fact, the association of VIFs with cortical arrays of F-actin stress fibers occurs at multiple length scales. For example, VIFs run through and frequently appear to interconnect with adjacent stress fibers, forming meshwork that surrounds them. These organizational states are consistent with the formation of an IPN. We show that this IPN structure has important functional consequences in cells and can result in enhanced contractile forces. Moreover, our results indicate that specific associations exist between actin and vimentin proteins in the cytoplasmic environment, which may facilitate the formation of an IPN; the results also show that the VIF network can influence the diffusive behavior of actin monomers, which may, in turn, have downstream effects on other actin-driven processes. Thus, vimentin has a far more comprehensive role in cellular function than previously thought. These findings confirm the importance of the interplay between VIFs and F-actin, especially as it relates to the formation of IPNs and their consequences on the contractile nature of cells.

## Results

### Vimentin intermediate filaments are present in the cortical region of MEFs

To investigate the details of the structural relationships between VIFs and F-actin, we image their organizational states in MEFs using structured illumination microscopy (SIM). Overall, F-actin networks are clearly defined along the periphery of the cell (Figure 1A). By contrast, networks of VIFs are more abundant deeper in the core of the cytoplasm (Figure 1B). However, in regions of the cortical cytoplasm, including protrusions, there exist both VIFs and F-actin, some of which are in the form of bundles or stress fibers (Figure 1C). Since the cytoplasm is thinly spread near the outer edge of the cell, the overlapping signals of VIFs and F-action must reflect the close proximity of the two types of filamentous cytoskeletal proteins.

**Figure 1.**
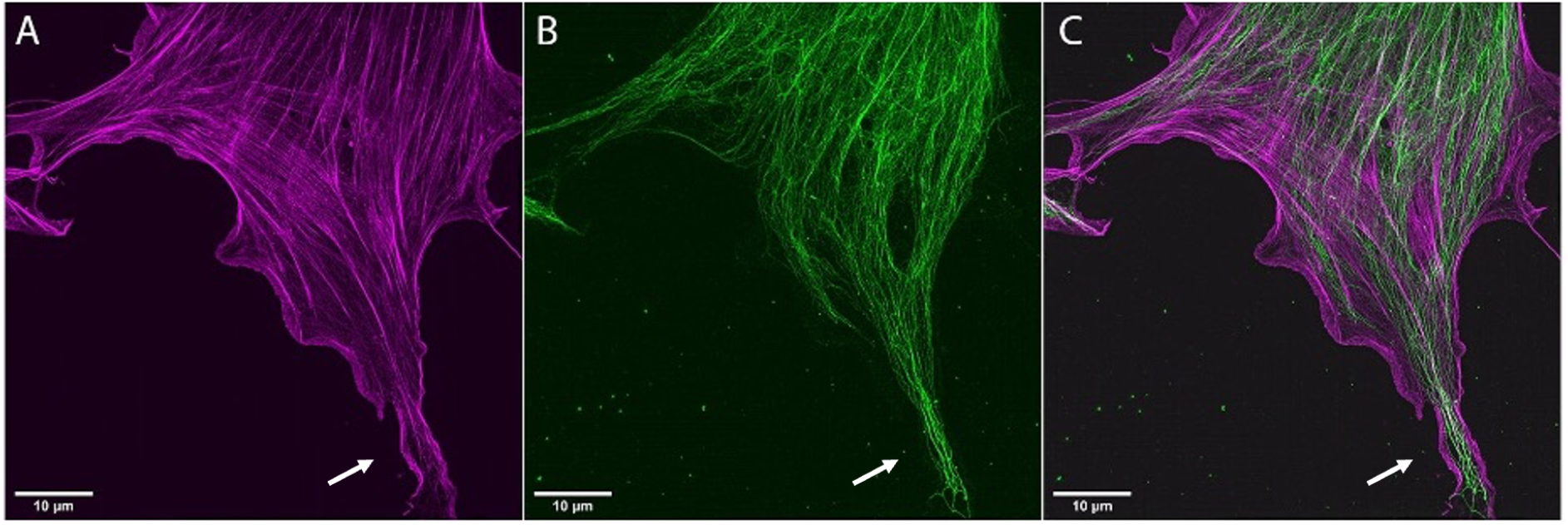
Mouse embryonic fibroblasts (MEFs) typically have an actin-rich periphery and cortex and a cytoplasmic core that contains more concentrated VIFs. F-Actin is indicated in magenta, VIFs in green. White arrows point to the protrusions. Scale bar represents 10 μm.

To better understand the relationship between the structures of F-actin and VIFs, we examine the basal side of the cell cortex in the region of cell-substrate adhesion with actin stress fibers. This reveals several distinct patterns that clearly suggest the formation of interpenetrating or interacting networks of these two cytoskeletal components. In some regions, there are distinct parallel arrays of closely spaced stress fibers and VIFs (Figure 2A). Most stress fibers are arrayed in well-defined, relatively straight tracks, whereas the VIFs are frequently in less well-oriented networks filling the space between stress fibers. Some stress fibers are not obviously associated with VIFs (Figure 2A). In other regions, we see thicker arrays of VIF, likely representing bundles, that run roughly perpendicular to the stress fibers, with some looser arrays of VIFs appearing to wrap around the stress fibers. This latter arrangement suggests that VIFs can form bridge-like structures between neighboring stress fibers (Figure 2B). The existence of such arrangements suggests that VIFs may modulate longer-range physical interactions between neighboring stress fibers. We also observe some sparser arrays of VIFs interlaced between or interconnecting stress fibers, forming a woven or interlaced structure (Figure 2C). Here, the VIFs alone are primarily oriented in a cross-hatched network fashion, and the stress fibers, all oriented in roughly the same direction, are woven through the VIF meshwork. Additionally, we find regions of co-aligned and co-located VIF fibers and stress fibers (Figure 2D). In these regions, all of the filaments are long and fully entangled, in each case resembling an interpenetrating network of polymers. This diversity of composite structures suggests that structure-coupling between F-actin and VIFs may be adaptable for different functions.

**Figure 2.**
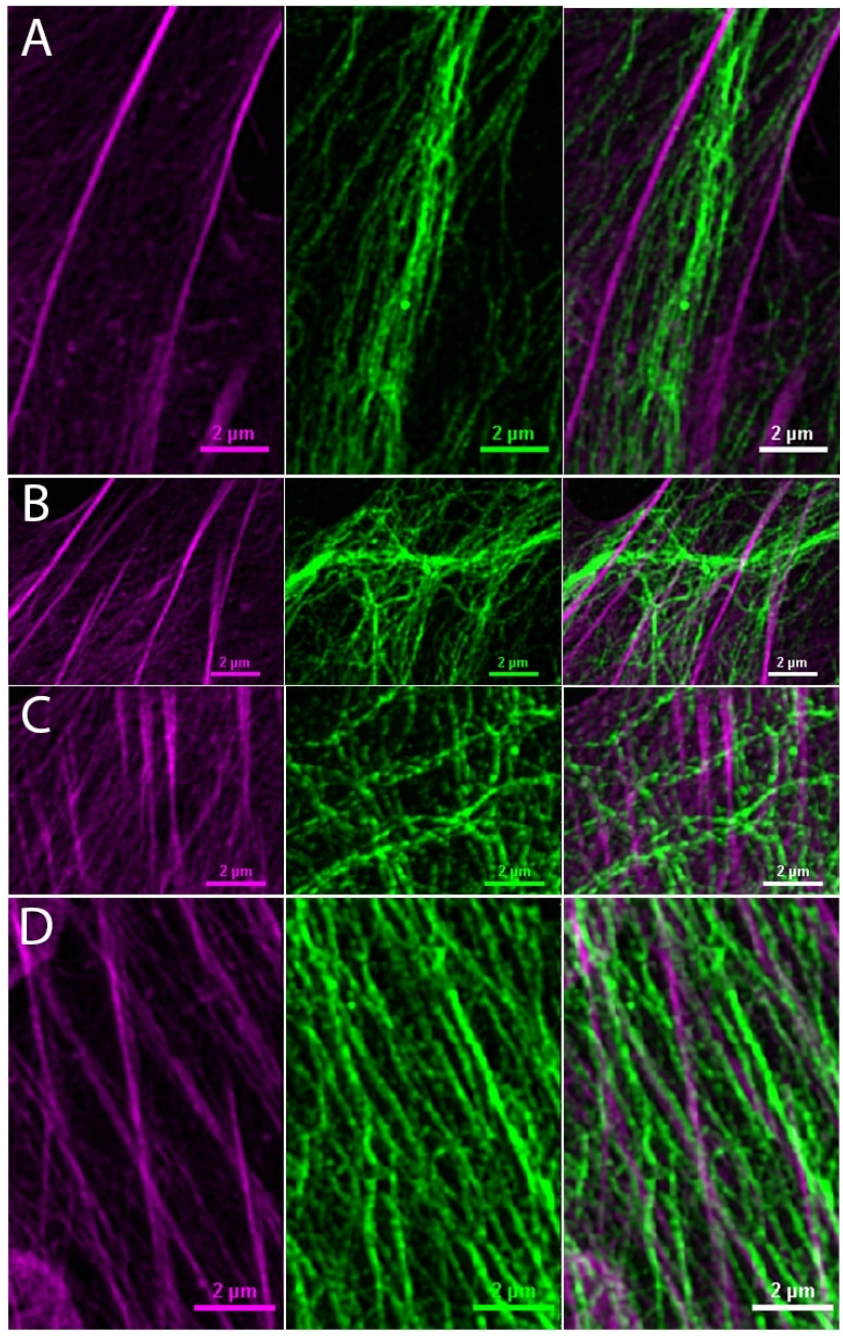
F-actin containing stress fibers and VIFs are in close proximity to the cell surface in the region of cell-substrate adhesion, as imaged by SIM: A) parallel arrays, B) bridging, C) interlaced, and D) close parallel arrays. The leftmost panels show F-actin, indicated in magenta. The middle panels show VIFs in green. The rightmost panels are merged images. Scale bar represents 2 μm.

### Cryo-electron tomography reveals a VIF and F-actin mixed polymer network in stress fibers

To accurately acquire a higher resolution view of the relationship between the structures of VIFs and F-actin within the cell cortex, we use cryo-electron tomography (cryo-ET). For this purpose, live MEFs grown on electron microscope grids are vitrified for structural studies, thereby maintaining their 3D structural organization (Medalia et al., 2002). To optimize the imaging of both VIF and F-actin in and around stress fibers, electron micrographs are taken in regions of protrusions in MEFs expressing emerald-vimentin. As observed by SIM in Figure 1, such protrusions invariably contain stress fibers, enabling us to carry out correlative light and cryo-ET of VIFs and F-actin (Figures 3A, B). We acquire tilt-series images and reconstruct the respective tomograms. The X-Y slices through the tomograms reveal VIFs in close proximity to the F-actin in stress fibers (Figure 3B). The F-actin fibers (red arrowheads) are long and straight 8-nm-diameter filaments, consistent with their long persistence length (Gittes et al., 1993). However, single VIFs (green arrowheads) are 11 nm in diameter and are bent and wavy, consistent with their much shorter persistence length (Goldie et al., 2007) (Figure 3B). In this and numerous other regions of the cell cortex (not shown) both VIFs and F-actin are aligned with their long axes approximately parallel, an arrangement reminiscent of the co-aligned fiber bundles observed with SIM (Figure 2).

**Figure 3.**
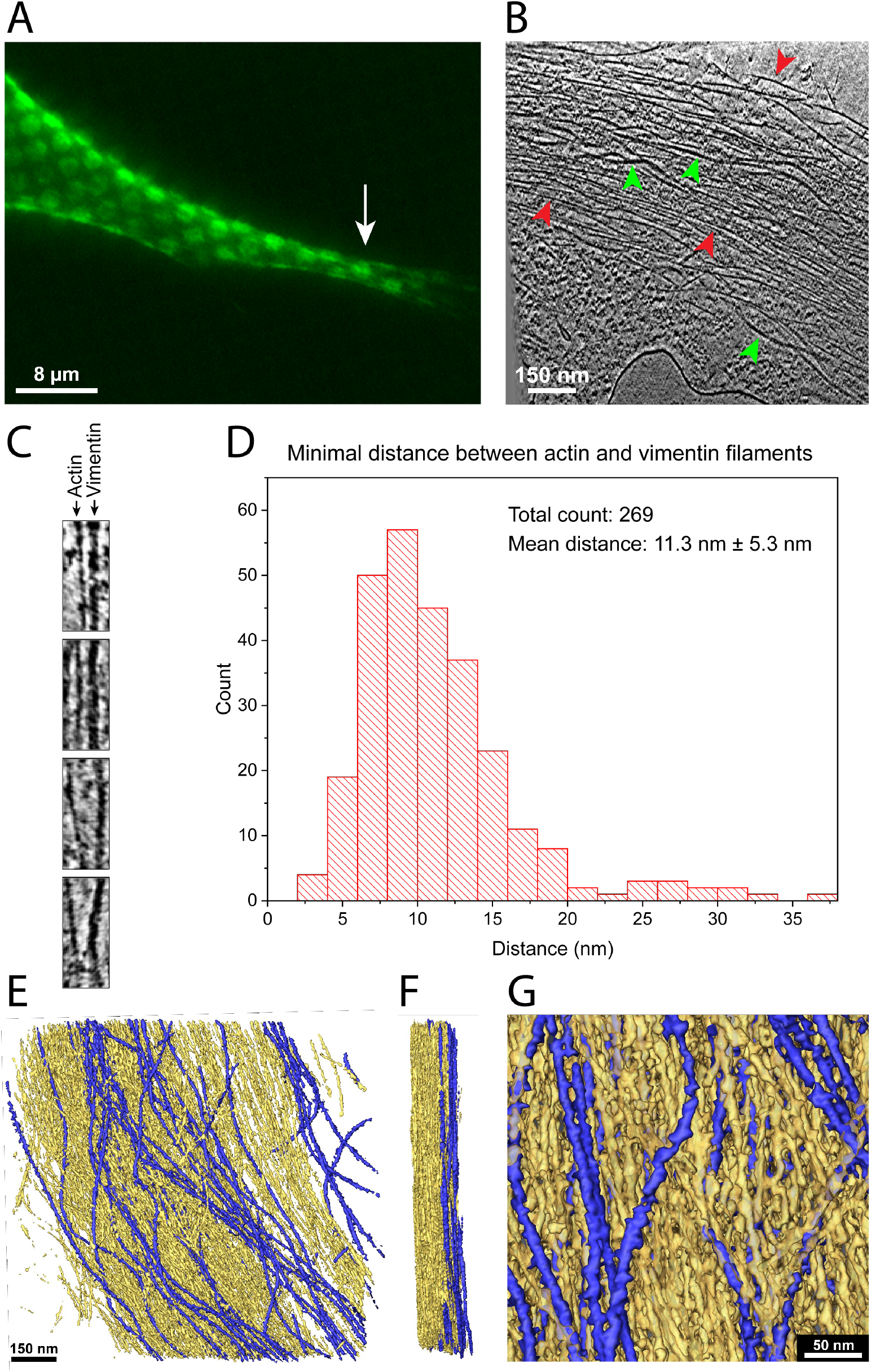
F-Actin-VIF co-localization in stress fibers is revealed by cryo-ET. MEFs expressing emerald-vimentin are grown on electron microscopy grids and imaged by fluorescence as well as electron microscopy. A) Fluorescence image of a MEF cell protrusion. Emerald-vimentin is shown in green. The white arrow indicates the position where a cryo-tomogram is acquired. The grid pattern seen in the fluorescent signal is due to the EM grid substrate. B) A 1.4 nm thick slice through a tomogram of a stress fiber, acquired at the cell surface facing the substrate to which the cell is attached. This region is the same as that indicated by the white arrow in A. Individual VIFs (green arrowheads) can be detected within the F-actin (red arrowheads) bundle. C) Representative images showing F-actin and VIFs in close proximity. F-actin is shown on the left, while VIFs are shown on the right. Scale bar: 50 nm. D) The minimal distances between F-actin and VIFs are calculated from cryo-tomograms of stress fibers in different cells (n=3). A total of 269 distance measurements between VIFs and F-actin filaments are made. The nearest neighbors are found within 11.3 ± 5.3 nm. E) Surface rendering views of the tomogram shown in B with VIFs (blue) and F-actin (yellow). F) Perpendicular view of the surface rendering seen in E shows that VIFs are primarily located at the basal (attachment) surface of the cell. Scale bar as in E. The thickness of the entire stress fiber in E and F is 236 nm. G) A detailed view of the F-actin-VIF composite network shows the close proximity of the two cytoskeletal elements.

To quantify the distance between VIF and F-actin filaments, measurements are made from three tomograms acquired from three different cells. Details of tomogram sections, showing representative examples of neighboring VIFs and F-actin from which measurements are made, are shown in Figure 3D. Measurements are made at points where a VIF lies closest to a parallel Factin filament. Specifically, the center-to-center distance between the two types of filaments is determined, and then the average radii of VIFs and F-actin are subtracted; the distance, therefore, represents only the space between the two types of filaments. The distance between VIF-F-actin is very short, averaging 11.3 ± 5.3 nm for 269 measurements (Figure 3C).

We also create 3D maps of surface renderings of regions of the cell cortex containing VIF and F-actin (Figure 3E-G). Interestingly, we find that in these regions, VIFs are mostly constrained to the basal side of the stress fiber (right side, Figure 3F). In close-up views from the basal surface of MEFs, the long axes of VIF filaments (blue) and F-actin (yellow) are mostly aligned (Figure 3E). Frequently the VIFs are either seen weaving in and out of the F-actin bundles or wrapping around them, while other regions are rich in F-actin alone. Such weaving can be clearly seen in Figure 3G. Importantly, it appears that the F-actin-rich stress fibers contain VIFs.

### VIFs impact F-actin functions in cell contractility

Since there is a close structural relationship between VIFs and F-actin in stress fibers, we investigate whether cell contractility is affected by mechanical interactions between these two cytoskeletal networks. To determine the role of VIFs, we use traction force microscopy (TFM) to compare cell contractility in vimentin-knockout (Vim-/-) and wild-type (WT) MEFs. We seed cells onto soft polyacrylamide gels with a layer of tracer particles embedded in the gel near its surface and image the positions of the tracer particles. By comparing their positions with and without cells present, we determine the displacements induced by the cells (Butler et al., 2002). We use these displacements to calculate the forces exerted by the cells on their underlying soft substrate. To ensure that the calculated forces can be attributed to single cells, we use sparsely seeded cells with the average distance between neighboring cells larger than the range of the strain fields from individual cells. Traction forces tend to be concentrated at the ends of cells, where the majority of the focal adhesions are located. To obtain a representative measure of the distribution of forces over the area of the cell, we determine the contractile moment, which is a scalar measure of contractile strength that assumes that the cell applies equal and opposite point forces whose separation is related to its size (Butler *et al*., 2002). Using this method, we have shown that WT MEFs are, on average, ~46% more contractile than Vim-/- cells (Vahabikashi et al., 2019).

To more precisely probe the independent contributions of F-actin and VIFs to contractility, we combine TFM with a transient stretch of the polyacrylamide substrate. We adjust the magnitude of the stretch such that it is insufficient to rupture the F-actin network but rather induces rapid disassembly and fluidization of the F-actin network into actin monomers (G-actin) mediated in part by the activity of actin severing protein cofilin (Krishnan et al., 2009; Lan et al., 2018; Trepat et al., 2007), followed by slow reassembly and resolidification of the F-actin network mediated in part by the activity of actin regulating protein zyxin (Rosner et al., 2017). Under these conditions of relatively little stretching, the highly extensible VIFs remain in the purely elastic regime (Janmey et al., 1991; Qin et al., 2009) and should not be affected by the stretch. To apply the stretch to the cells, we use a cylindrically-shaped plastic indenter attached to a motorized arm. The indenter contacts the gel substrate in a 3 mm ring centered around the cell of interest and compresses the gel to introduce a 10% stretch at the center for 3 seconds. A schematic of the setup is presented in Figure 4A. To study the dynamics of recovery, we use TFM to track the contractility of the cells over 10 minutes as the cell reassembles the fluidized actin monomers back into F-actin filaments and stress fibers (Krishnan *et al*., 2009). The actin fluidization causes an immediate drop in cell contractility from the baseline state, and the contractility recovers as the cell rebuilds its contractile framework, as shown by the representative data for the recovery of the contractility for both WT MEFs and Vim-/- MEFs in Figure 4B.

**Figure 4.**
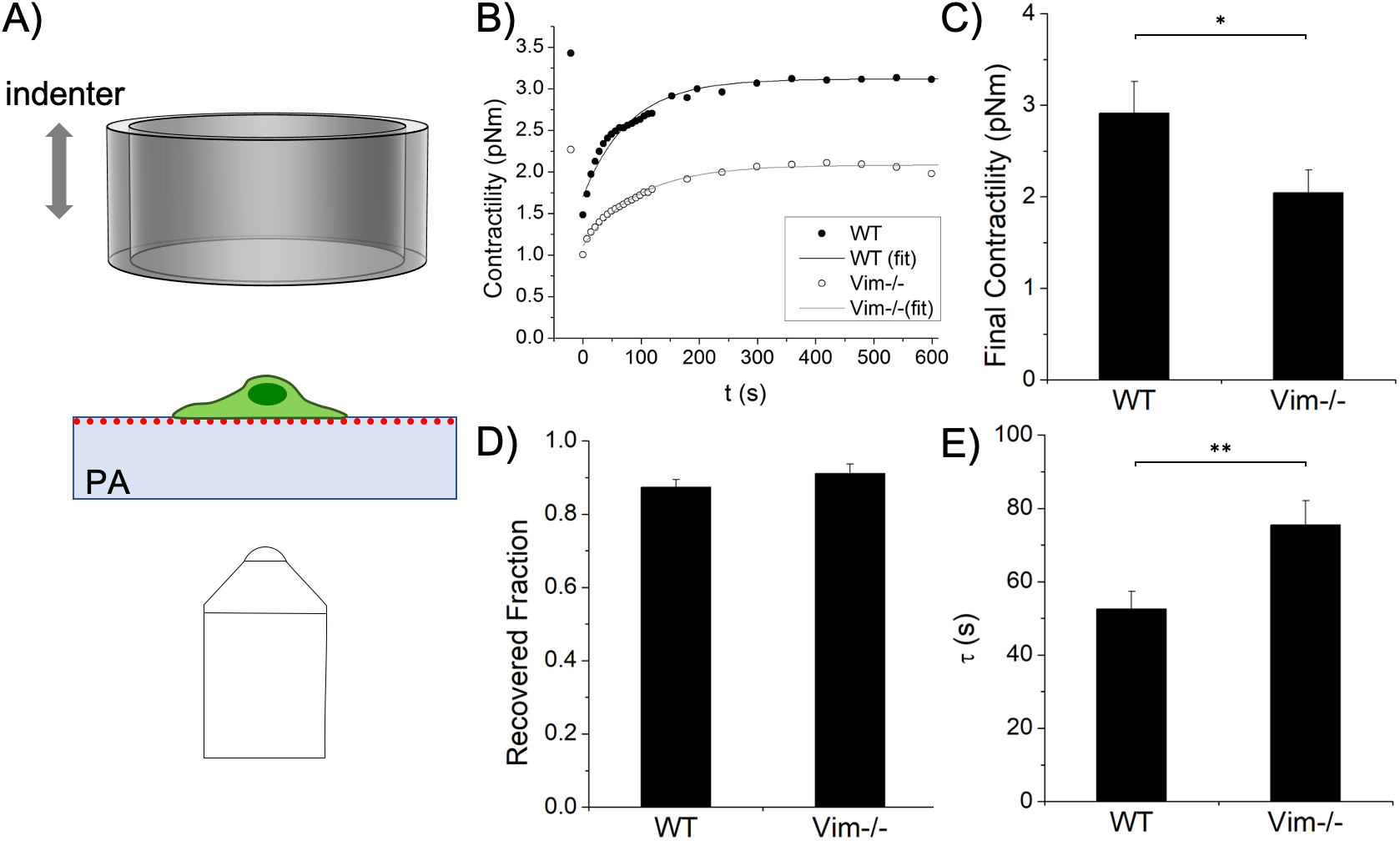
Transiently stretching cells fluidizes the F-actin cytoskeleton, but cell contractility recovers within several minutes coincident with its reassembly. A) Schematic of the stretching setup. Cells are grown on a soft polyacrylamide gel embedded with beads. They are stretched isotropically by a large cylindrical indenter centered around the cell. B) Representative recovery curves for WT and Vim-/- MEFs and fitting curves used to quantify the recovery dynamics. Vim-/- cells are less contractile after recovery (C; p<0.05) but both cell types recover about the same fraction (D). E) Vim-/- cells take longer to recover (p<0.01). All results are plotted as mean+/-SEM.

At the end of the recovery period following stretching, WT cells remain ~43% more contractile than Vim-/- cells (Figure 4C). This difference is similar to that observed in unstretched cells; thus, decreased contractility is a characteristic of the Vim-/- MEFs. Furthermore, all the cells exhibit a plateau at a similar fraction of their initial contractility, ~90%, regardless of vimentin expression (Figure 4D). This shows that the cells experience little or no permanent damage due to the stretch; within a short time, the cells are able to recover most of the contractility they generate upon their normal attachment and spreading on the substrate.

To quantify the recovery dynamics, we fit the recovery curve with an exponential function and determine a time constant τ. Cells lacking vimentin are slower to recover, taking 75.4s on average; by contrast, WT cells recover in 52.5s on average (p<0.01, Figure 4E). The Vim-/- cells take longer to reach their steady-state contractility, implying that these cells build their actomyosin system more slowly. Moreover, these results suggest that the assembly of F-actin based contractile networks can be regulated by VIFs in mesenchymal cells such as fibroblasts.

### Vimentin decreases G-actin diffusion

To further explore the functional relationship between VIFs and F-actin, we investigate the effect of VIFs on the diffusive behavior of G-actin using fluorescence recovery after photobleaching (FRAP). We transfect WT and Vim-/- MEFs with EGFP-actin and confirm that the stress fibers in Vim-/- cells appear normal. To make the FRAP measurements, we briefly illuminate a small circular region that contains fluorescent stress fibers with an intense laser to locally photobleach the actin, and then measure the recovery of the fluorescence over time (Figure 5A). As GFP tagged G-actin diffuses back into the bleached region, the intensity recovers in an exponential manner until it reaches a plateau. To facilitate comparison between cells, the time dependence of the intensity is normalized to its initial value. The fluorescence intensity drops by ~ 40% during the bleaching step for both WT and Vim-/- MEFs and the recovery plateaus at ~ 80%, as shown for a WT (filled circles) and a Vim-/- MEF (open circles) in Figure 5B. Some of the actin monomers (G-actin) that are the subunits of F-actin remain bound on the timescale of the experiment (Ehrlicher et al., 2015), and this results in a fraction of fluorescence that cannot be recovered, designated as the immobile fraction. We fit the recorded intensities to an exponential I(t) = C - A * exp(-t/τ), where C is the immobile fraction and τ is the recovery time constant.

**Figure 5.**
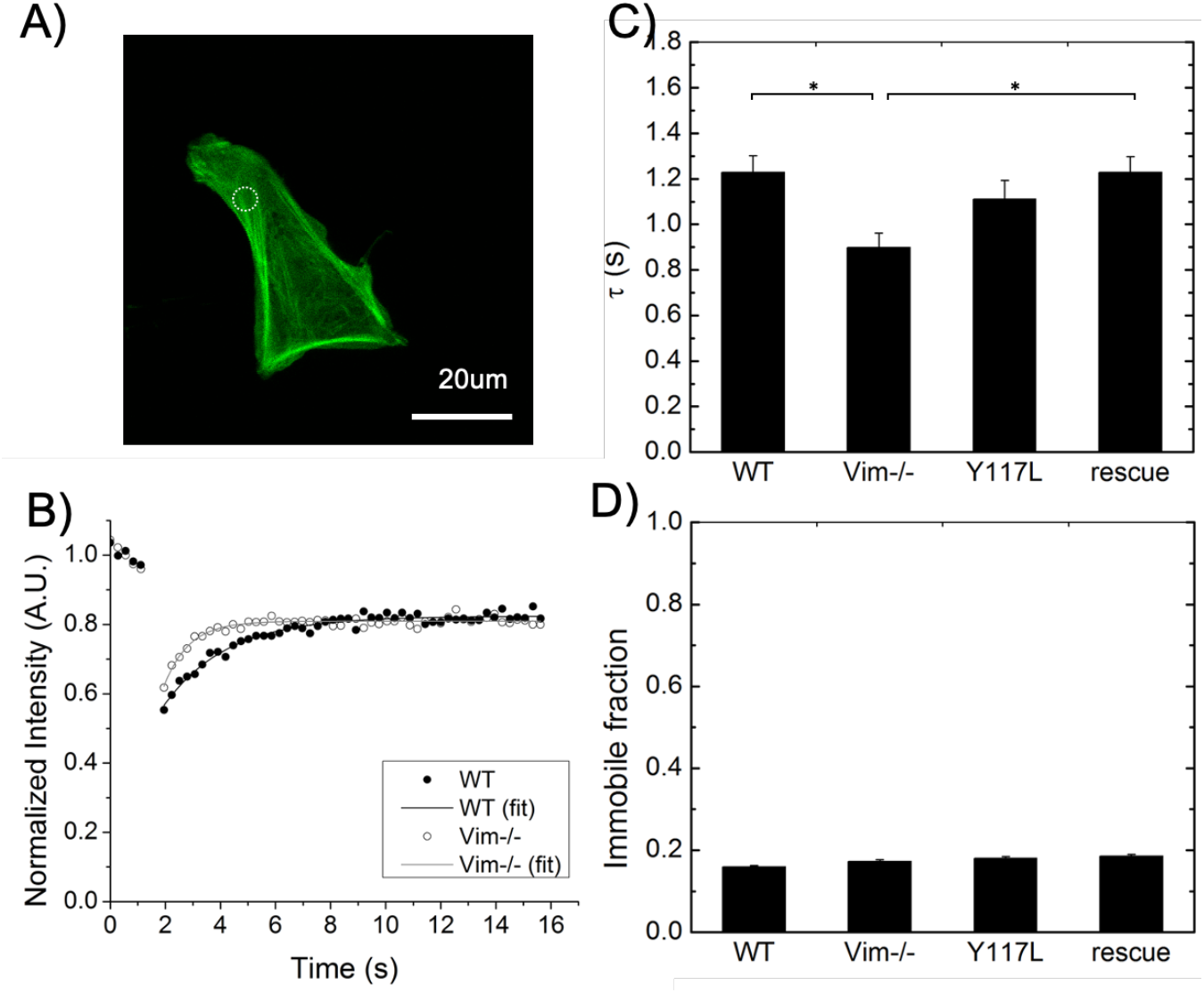
Fluorescence Recovery After Photobleaching (FRAP) using EGFP-tagged actin shows that in the presence of VIF, G-actin diffusion-like motion decreases A) EGFP-actin (green) and bleach spot (white dashed circle) of a sample MEF. B) Representative recovery curves for WT and Vim-/- MEFs. C) The immobile fractions are the same for all MEFs studied. D) The fluorescent stress fibers in Vim-/- MEFs recover fluorescence faster, indicating faster G-actin diffusion-like motion. All results are plotted as mean+/-SD (n = ~35).

The WT MEFs recover their fluorescence with τ = 1.2 s, which is about 33% slower than the Vim-/- cells, which have τ = 0.9 s (p<0.05; Figure 5C). This suggests that the VIFs inhibit the motion of actin monomers. Based on these values of τ, an effective G-actin diffusion coefficient is calculated to be 1.9 μm^2^/s for WT MEFs and 2.5 μm^2^/s for Vim-/- MEFs; we emphasize that these coefficients are not due to thermal diffusion but are rather most likely driven by the random forces due to motor or enzymatic activity within the cell (Ming Guo, 2014). By contrast, no significant difference is observed in the immobile fractions of the two cell types; both have a C of about 0.17 (Figure 5D). This suggests that the fraction of actin in the polymerized state is not significantly affected by the presence of vimentin.

To determine whether the difference in the diffusion-like motion is due to the VIF network structure, we carry out FRAP analyses in Vim-/- MEFs that express only vimentin with a Y117L mutation, which allows lateral assembly of monomers into unit length filaments (ULF) but prevents the end-to-end annealing that forms long VIF (Brennich et al., 2019; Herrmann et al., 2009). The ULFs are about the same width as fully-assembled filaments but only ~65 nm long. We find that the immobile fraction does not change (Figure 5B) but the effective diffusion coefficient of G-actin in these cells is in between those observed in Vim-/- and in WT MEFs (Figure 5C), suggesting that the difference is partially an effect of direct protein interactions rather than solely the physical obstruction due to a complex VIF network structure. We also analyze Vim-/- MEFs rescued by transfection with vimentin cDNA, which assemble long, mature VIFs, but whose vimentin protein expression is only ~30% of that of the endogenous concentration in WT MEFs. The rescued cell line is able to fully reproduce the slower diffusion-like motion of G-actin (Figure 5C). These results imply that the filamentous form of vimentin is most effective at inhibiting G-actin motion, while providing further support that the soluble forms of the proteins, both G-actin and ULF, are able to interact with each other in cells.

### In vitro reconstituted networks form IPN without the presence of other cellular components

Our findings that VIFs and F-actin form composite structures comprised of IPNs in the cortical region of cells and that VIFs can inhibit G-actin motion suggest that there may be a direct structural coupling between these two cytoskeletal elements. This possibility is further supported by their very close proximity within stress fibers (Figure 3). This close physical association suggests either the existence of specific crosslinking molecules or direct interactions between VIFs and F-actin. The only well-defined VIF-F-actin cross linking protein is plectin, which is comprised of >4000 amino acids with a molecular weight of ~500 kDa. Plectin is thought to form dimers consisting of a 200 nm long coiled-coil domain flanked at each end with large globular domains (Wiche, 1998). It is therefore unlikely that plectin could fit into the space separating VIF and F-actin in stress fibers. Another possibility is that these two cytoskeletal polymers can bind to each other directly (Duarte et al., 2019; Esue et al., 2006; Kolsch et al., 2009), which is consistent with our FRAP results. However, given their physical and chemical properties, it appears unlikely that these two cytoskeletal polymers would form mixed complexes as they are both highly negatively charged and therefore would be unlikely to co-assemble in vitro. We therefore explored the structural properties of a model in vitro system consisting of reconstituted purified VIFs and F-actin at approximately physiological concentrations. The vimentin we use is extracted and enriched as polymerized VIFs from WT MEFs rather than expressed in bacteria; thus, reflecting a more physiological state, and is more likely to retain post-translational modifications. To assemble the model system, we develop a buffer in which purified preparations of both proteins assemble *in vitro*. The reconstituted networks are imaged using a confocal microscope. The vimentin preparation assembles into a network of VIFs that are several microns long and very flexible in appearance with a persistence length that is less than a micron (Figure 6A). Actin also assembles in the same buffer and forms a network of relatively straight filaments, consistent with the expected persistence length of ~17 microns (Figure 6B).

**Figure 6.**
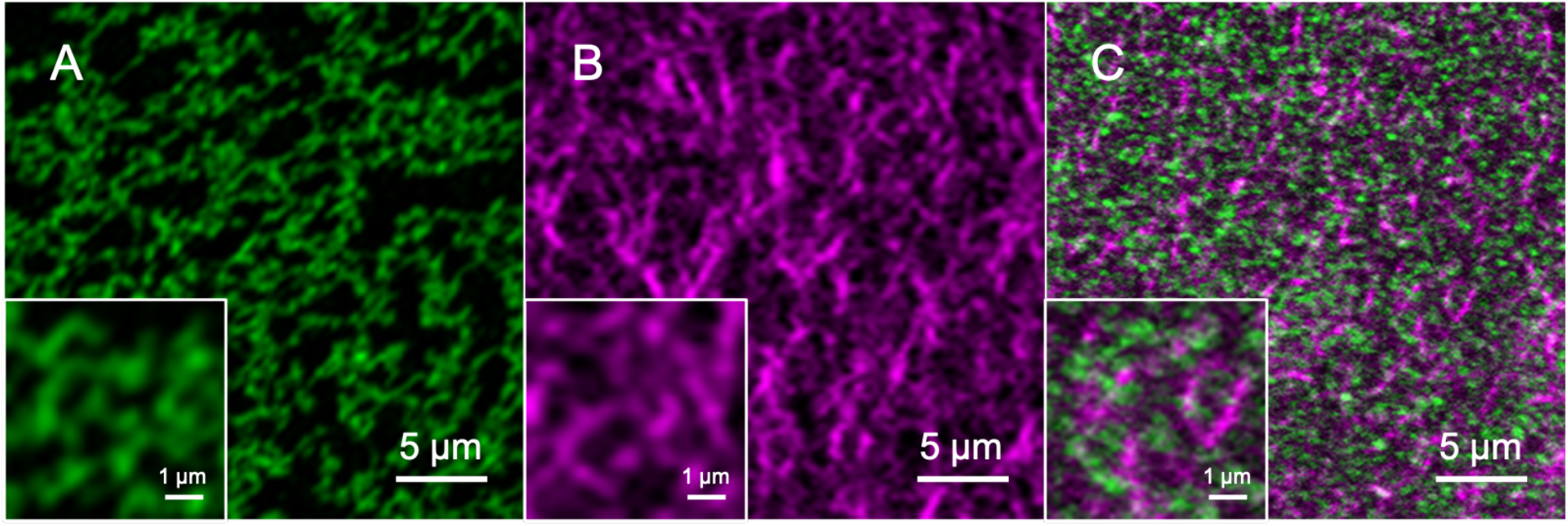
Confocal fluorescence images of reconstituted networks of A) VIFs (green), B) F-actin (magenta), and C) a mixture of VIFs and F-actin. The insets show enlarged views of the networks. All three samples use the same buffer conditions for assembly.

When both F-actin and VIFs are polymerized together, the proteins self-assemble into randomly oriented filaments (Figure 6C). Importantly, no large-scale phase separation is detected; instead, we only observe interpenetrating networks of the two proteins. However, we do not observe lateral associations between individual filaments over long distances, as observed in cells. In this reconstituted system, the divalent cations, which are necessary for actin polymerization, may form transient crosslinks between the two types of proteins and thus may help facilitate IPN assembly. Within this IPN, the highly flexible filaments of the VIF network seem to fill in the pore spaces of the much more rigid filaments of the F-actin network. Since the F-actin fibers in a cell are always under more tension due to contributions from motor proteins and numerous crosslinking proteins, this reconstituted system cannot fully reflect the cytoplasmic structures *in situ*, as observed by SIM (Figures 1–2) and Cryo-ET (Figure 3); nevertheless, these results confirm that Factin and VIFs can form interpenetrating networks in the absence of other cell components.

## Discussion

The results presented in this paper provide strong evidence that VIFs exist in significant quantities in the cell cortex. They form an interpenetrating network with the F-actin network resulting in strong elastic interactions between the two networks; in addition, they exist in very close proximity with the F-actin stress fibers. This is contrary to the widely prevalent belief that VIFs are compartmentalized in the bulk cytoplasm and that F-actin and associated proteins are the proteins that define the properties of the cell cortex. Stress fibers are typically thought to consist of bundles of F-actin filaments and their closely associated proteins such as myosin (Livne and Geiger, 2016). However, the VIF network forms distinct arrangements with stress fibers over microns-long regions. Strikingly, high-resolution analyses of peripheral stress fibers by SIM and Cryo-ET reveals that VIFs are integral components of the stress fibers themselves. Indeed, individual VIFs course through and around the actin bundles. In fact, the average minimal interfilament spacing between individual VIF and F-actin filaments is smaller than between the neighboring F-actin (~22 nm) in stress fibers (Martins et al., 2021), which is determined by the size of crosslinkers such as alpha-actinin or full-length vinculin (Boujemaa-Paterski et al., 2020). Thus, any interactions that exist between the two filaments could be direct or could be mediated by one or more very small unknown crosslinkers. The finding that VIFs are woven into F-actin stress fibers suggests that they may become incorporated at the time of stress fiber formation, and not as a secondary structure that is added later. Indeed, confocal images show that reconstituted Factin and native, mammalian cell-expressed VIFs that are polymerized simultaneously *in vitro* form an interpenetrating network even without the help of binding proteins; this is also consistent with electron micrographs of reconstituted composite networks when both proteins are bacterially expressed (Esue *et al*., 2006). This suggests that our observations of the interpenetrating network structures in cells, using SIM and Cryo-ET, reflect fundamental properties of VIFs and F-actin. However, this reconstituted system cannot fully replicate the structures we observe in cells, indicating that other factors are necessary. This will be an important area of further study using reconstituted networks of increasing complexity.

The variety of structures and the range of length scales over which VIFs and F-actin associate suggest that these composite interpenetrating networks may be widely involved in cell functions attributed to the cell cortex. We show that the integrated F-actin and VIF structures have important functional consequences for cell contractility: the presence of VIFs enhances both the magnitude and rate of actin-generated contractility (Jiu et al., 2015; Patteson et al., 2019; Vahabikashi *et al*., 2019). Although the VIF network is not itself a contractile system, our observations suggest that this structural polymer nevertheless plays an essential supporting role in cellular force generation and its consequences. For example, the different rates of recovery from stretching may help explain why cells lacking vimentin exhibit slower migration speeds, whereas cells that express vimentin during the EMT or cancer metastasis are more migratory (Eckes *et al*., 1998). Migration requires cells to pull themselves along a substrate in a directional manner; in cells that lack vimentin, the effects of slower force generation and lower contractility will combine to result in impaired migration. This is consistent with observations that vimentin-deficient cells and the mice from which they are derived are more prone to damage upon experiencing tensile stress (Eckes et al., 2000; Eckes *et al*., 1998) or wounds (Cheng et al., 2016), or during migration through small pores (Patteson *et al*., 2019); while we do not stress cells to the point of rupture in this study, the slower recovery of Vim-/- cells will lead to a reduced resilience against stress. In addition, these results hint that VIF may be able to regulate the assembly of actin/myosin networks.

While the filamentous forms of vimentin and actin clearly work synergistically, it is also important to consider interactions between other forms of the proteins. In particular, there is evidence for large pools of G-actin in the cytoplasm, whose concentration drives the local polymerization and dissociation of F-actin (Atkinson et al., 2004). By contrast, vimentin is mainly found in the form of assembled VIFs (Herrmann et al., 1996), while its soluble pool is much less. Assuming a homogenous distribution of proteins, a rough calculation of the volume taken up by the vimentin and actin monomers estimates each at about 1% volume fraction in the cytoplasm (Koffer et al., 1983). However, in their fully assembled forms, each protein network has a much higher effective volume fraction, of 25% or greater (Supplemental Materials). Thus, the fact that G-actin moves more slowly in WT cells than in Vim-/- cells could be due to obstruction of G-actin by the VIF network, similar to the observation that cells with VIF exhibit significantly reduced organelle motion (Guo et al., 2013). However, given the tiny size of G-actin compared with the VIF-network mesh size, it may be that some transient attractive interactions or binding between the two proteins also contributes to the reduced motion. Our FRAP results in MEFs with the Y117L mutation support this, since even the unpolymerized vimentin ULFs are able to slightly influence the fluorescence recovery time. Since EGFP is a relatively large molecule, about half the size of G-actin itself, these measurements may underestimate the effective diffusion coefficient compared with that of unlabeled G-actin. However, these results are still roughly consistent with other reports of the G-actin diffusion coefficient in cells, which ranges as low as 2 μm^2^/s (McGrath et al., 1998). Furthermore, although the motion we observe is actively driven, the coefficients we obtain are orders of magnitude lower than that expected from a cytosolic viscosity similar to that of water. Thus, there may be environmental factors that slow the G-actin motion, one of which could be VIFs. The fact that vimentin in either its fully polymerized or partially assembled states can affect the motion of G-actin could be an important means by which cells can locally tune their mechanics and G-actin distribution, with possible downstream effects on actin polymerization and other dynamic processes.

Taken together, these results suggest that the structure and the mechanical behavior of the cortical region are both consistent with interpenetrating networks of crosslinked hydrogels. For example, when an ionically-crosslinked gel is combined with a covalently crosslinked gel, the hybrid gel would exhibit extreme toughness due to load sharing by the two networks (Sun et al., 2012). The weaker ionic crosslinks can dissociate to dissipate force but soon reform once the load is removed, thereby protecting the other network from rupture. Furthermore, inter-network bonds can promote self-healing by preserving some memory of the initial configuration. In cells, VIFs form an ionically crosslinked gel (Lin et al., 2010) and F-actin has many crosslinking proteins, some of which bind for long time scales and others of which are weaker. When both cytoskeletal networks are present, the cells are tougher (Hu et al., 2019) and more motile (Mendez et al., 2014), which is correlated with increase in cell stiffness. As a cell is stretched, the highly extensile VIF network may help to dissipate some of the stress. Since actin disassembles rather than ruptures, the VIF are unlikely to have a direct role in fluidization. However, during recovery, the VIF network could function to maintain a locally high G-actin concentration through physical associations. This would help recover the original stress fiber structure and result in a quicker return to the baseline state as is observed.

Overall, these results highlight that VIFs play a much broader role in cellular mechanics than previously thought. As an integral component of stress fibers, VIFs can contribute not only to the resilience of a cell, but also to dynamic processes such as contractility. Moreover, since VIF or ULF can modify the behavior of G-actin subunits, they may also indirectly influence many other processes that are driven by actin. There is no doubt that VIFs and F-actin are both significant mechanical contributors in cells; however, the results presented here strongly support that their contributions are strongly correlated.

## Acknowledgments

We gratefully acknowledge Dr. Bomi Gweon for insightful discussions and assistance with traction force measurements. This work was supported in part by NIH Grants 2P01GM096971 awarded to RDG and DW, and 1R01HL148152 and 1U01CA202123 awarded to JJF. DW also acknowledges support from the Harvard Materials Research Science and Engineering Center (MRSEC) Grants DMR-1420570 and DMR-2011754. O.M. acknowledges support by The Swiss National Foundation, grant no. 31003A_179418. M.W. was funded by the Forschungskredit fellowship from the University of Zurich. Y.S. was supported by the NSF-Simons Center for Mathematical and Statistical Analysis of Biology at Harvard (NSF award no. 1764269) and the Harvard Quantitative Biology Initiative.

## Materials and Methods

### Cell culture

Wild-type mouse embryo fibroblasts (WT mEFs) and vimentin-null mouse embryo fibroblasts (Vim-/- mEFs) are kindly provided by J. Eriksson (Abo Akademi University, Turku, Finland) and are maintained in DMEM with 25 mM HEPES and sodium pyruvate (Life Technologies; Grand Island, NY) supplemented by 10% fetal bovine serum, 1% penicillin streptomycin, and nonessential amino acids. All cell cultures are maintained at 37°C and 5% CO2. The vimentin-null mEFs expressing vimentin are created by PCR amplification of the vimentin coding sequence using CloneAmp polymerase (Clontech) from pcDNA4-vimentin (provided by J. Eriksson) and the coding sequence for Vimentin Y117L is amplified from pmCherry-C1-Vim Y117L (provided by H. Herrmann) using the primers ggcgccggccggatccATGTCCACCAGGTCCGTGTCC and actgtgctggcgaaTTATTCAAGGTCATCGTGATGCTGAG. The PCR product is purified from an agarose gel and inserted into pBABE-hygro (pBABE-hygro is a gift from Hartmut Land & Jay Morgenstern & Bob Weinberg (Addgene plasmid #1765; http://n2t.net/addgene:1765; RRID:Addgene_1765) cut with BamHI and EcoRI using In-Fusion (Clontech). Virus is produced by transfection of 293FT cells with pBABE-vimentin and pCL-Eco using Xfect transfection reagent (Clontech) and collection of supernatants 48 and 72 hours post transfection. The pooled virus supernatants are diluted in fresh complete medium and brought to 8 μg/ml polybrene prior to addition to vimentin-null mEFs. The virus supernatant is removed after 6 hours and replaced with fresh media. 24 hours after the first application of virus supernatant, the process is repeated. 48 hours following the second application of virus, the medium is replaced with fresh complete medium containing 200 μg/ml hygromycin. The selection medium is changed every two days for 7 days with the culture passaged as needed.

### Sample preparation for cryo-ET

MEFs expressing emerald-Vimentin are cultured on glow-discharged holey carbon coated EM grids (Au R2/1, 200 mesh, Quantifoil) for 16 h at 37°C in a humidified CO2 incubator. Cells are rinsed in PBS, fixed with 4 % paraformaldehyde (PFA) for 5 min and washed again in PBS. The cells are imaged by fluorescence microscopy (Leica DMI 4000B, Leica) using a 63x objective. Next, the grids are vitrified in liquid ethane after the addition of 10 nm gold fiducial markers (Aurion, Netherlands).

### Cryo-electron tomography-data acquisition and image processing

Tilt series are acquired using a Titan Krios electron microscope (ThermoFisher) operated at 300 KeV and equipped with a K2 Summit direct electron detector (Gatan) mounted on a postcolumn energy filter (Gatan). Ten tilt series are acquired in a zero-loss energy mode with a 20 eV slit. The data are acquired at a magnification of 42,000 x resulting in a pixel size of 0.17 nm in super-resolution mode and a defocus of −3 μm. A bidirectional tilt scheme with a tilt range of ±60° and an increment of 3° is chosen, corresponding to 41 projections per tilt series and a total cumulative electron dose of ~55 e/Å^2^. SerialEM 3.5.8 (Mastronarde, 2005) is used for data acquisition. A correlative light and electron microscopy (CLEM) approach is used, namely, tilt series are acquired at positions where vimentin IFs are identified in the fluorescence microscopy images.

The projection images are binned and subjected to motion correction using MotionCorr (Li et al., 2013), resulting in a final pixel size of 3.4 Å. Next, tomograms are reconstructed in a size of 1024×1024×512 voxels (final voxel size: 13.6 Å) using the TOM Toolbox (Nickell et al., 2005). Vimentin and actin filaments present in the tomograms are manually segmented using the Amira 5.6.0 software package (Thermo Fischer Scientific). This software is also used to analyze the distances between vimentin and actin filaments and for visualization purposes. In addition, OriginPro 2018 software (OriginLab Corporation) is used for distance measurement evaluation and visualization. The distance is measured from the center of vimentin to the center of actin and then the average radii of actin and vimentin are subtracted. The distance therefore represents only the space between the filaments.

### Structured illumination microscopy (SIM)

Mouse embryonic fibroblasts are seeded on #1.5 glass coverslips and fixed with 4% PFA for 10 min at RT. The fixed cells are permeabilized with 0.1% Triton-X 100 for 10min at RT and stained with anti-vimentin (1:200, Biolegend, CA, USA) for 30 min in PBS containing 5% normal goat serum (RT). This is followed by incubation with goat anti-chicken Alexa Fluor 488 (1:400, Invitrogen, CA) and Alexa Fluor 568 phalloidin (1:400, Invitrogen, CA) in PBS for 30 min. (RT). The coverslips containing the stained cells are mounted with ProLong Glass antifade mountant (Life Technologies, Carlsbad, CA, USA) on microscope slides. 3D SIM is carried out with a Nikon N-SIM Structured Illumination microscope system (Nikon N-SIM, Nikon, Tokyo, Japan) using an oil immersion objective lens (CFI SR Apochromat 100x, 1.49 NA, Nikon). For 3D SIM, 10 optical sections are imaged at 100 nm intervals in the periphery of the cell. Raw SIM images are reconstructed with the N-SIM module of Nikon Elements Advanced Research with the following parameters - Illumination contrast: 1.00; high-resolution noise suppression: 0.75; out-of-focus blur suppression: 0.25. Brightness and contrast are adjusted for image presentation.

### Reconstitution of purified F-actin and vimentin IF

We extract vimentin from mouse embryonic fibroblasts (MEFs), which are grown in dishes and washed 3 times with PBS. Lysis buffer (0.6 M KCl; 10 mM MgCl_2_; 1% TritonX-100; 1 mM PMSF) is added to the cells and the lysate is placed in a homogenizer for 5-10 min. DNaseI is added at a concentration of 1 mg/ml to the lysate and then centrifuged at 1600 g for 15 min at 4 °C. The pellet is washed 3 times (5 mM EDTA; 0.2 mM PMSF in PBS) and suspended in disassembly buffer (8 M urea; 5 mM NaPO4 pH 7.2; 1 mM PMSF; 0.2% mecaptoethanol) after which it is stirred for 45 min at RT. The suspension is centrifuged at 75000 rpm for 30 min at 20°C to clarify it. The supernatant is dialyzed overnight at room temperature against a large volume of buffer (0.1 mM 2-mercaptoethanol; 0.1mM PMSF in PBS). The dialysate is used for further experiments. This procedure for isolating and reassembling vimentin IF is modified from a previously published protocol (Zackroff and Goldman, 1979).

We mix dialyzed vimentin, rhodamine-labeled G-actin (AR05, Cytoskeleton Inc) and unlabeled G-actin (AKL99, Cytoskeleton Inc) successively into the assembly buffer and let them equilibrate at 37 °C for 1 hour. The assembly buffer is as follows: 10 mM Tris-HCl (pH 7.5), 2 mM MgCl_2_, 50 mM KCl, 1 mM ATP, 5 mM guanidine carbonate, 170 mM NaCl, and 1 mM DTT. We use glutaraldehyde (16220, Electron Microscopy Sciences) to fix filaments on a coverslip for 5 minutes and gently wash them using PBS buffer. To visualize VIFs, we stain them using a primary antibody (1:200, CPCA-Vim, Encor Biotechnology Inc) and a secondary antibody (1:400, A-11039, Thermo Fisher Scientific) successively with each staining for 45 minutes at room temperature followed by washing with PBS buffer. The visualization of F-actin does not require antibody staining, as pre-labeled G-actin is assembled together with unlabeled G-actin in the ratio of 1:4. We image the networks using a confocal microscope (Carl Zeiss, LSM 510).

### Cell Stretching and Traction Force Microscopy

Collagen-coated polyacrylamide gels with a Young’s modulus of 2.4 kPa are prepared in 35 mm glass bottom dishes (In Vitro Scientific/CellVis) (Butler *et al*., 2002). Gels intended for traction force microscopy are prepared with 0.5 μm red fluorescent tracer particles embedded near their surface. Cells are sparsely seeded on the gels in the presence of culture medium and allowed to grow for 24 hours before experiments are carried out.

The cells are stretched using an indenter ring with a circular cross section attached to an arm controlled by custom-written LabView code. When initiated, the indenter applies and holds a 10% strain around the selected cells for 3 seconds before being lifted back up.

To perform traction force microscopy, a Leica epifluorescence microscope is used to image the tracer particles and the cells throughout the stretch and recovery period. Several images are taken before stretching to establish a baseline and images are taken at designated intervals following the stretch to monitor recovery. At the end of the time, the cells are removed by trypsinization and a reference set of images without attached cells is taken. Substrate displacements are analyzed by comparing the bead images with and without cells using particle image velocimetry (PIV) in a custom MATLAB code. Traction forces are calculated by applying a Fourier transform to the displacement field (Butler *et al*., 2002). The contractile moment is determined as an average measure of contractile force for each individual cell.

### Fluorescence Recovery After Photobleaching (FRAP)

WT and Vim-/- MEFs are transfected with an EGFP-actin plasmid using Lipofectamine 2000 transfection agent (Invitrogen) and imaged on the third day. Fluorescence recovery after photobleaching (FRAP) is performed (Ehrlicher *et al*., 2015). Briefly, transfected cells are bleached for 1s using the FRAP module within the Leica SP5 confocal software, and monitored for 30 seconds, acquiring an image every 0.5s using a 63x/1.2NA water-immersion objective. The measured intensities are normalized to the pre-bleach intensities of the region of interest (ROI), and the recovery curve is normalized by a control ROI to account for sample bleaching during image acquisition. Since the brightness varies from cell to cell, we also normalize the intensities to pre-bleach levels. The intensity recovery is fit by I(t) = C - A * exp(-t/τ), where τ is the time constant and C is the immobile fraction.

