## Supplemental Information for "Vimentin Intermediate Filaments and Filamentous Actin Form Unexpected Interpenetrating Networks That Redefine the Cell Cortex"

**Estimation of volume fraction of F-actin and VIFs**

We estimate the effective volume fraction of F-actin and VIFs, which are both semiflexible polymers, in a cell using the worm-like chain model. We first calculate the filament length per unit of volume as $\rho \sim \frac{cN}{M_w}$ using the monomer molecular weight $M_w$, the protein concentration $c$, and the number of monomers per length of filament $N$. Then, we calculate the expected tube radius $R_e \sim 0.80\rho^{-3/5}L_p^{-1/5}$ using the persistence length of the polymer, $L_p$. We determine the effective total volume occupied by the filaments by assuming the total length of filaments in the cytoplasm is a cylinder of radius $R_e$. Finally, we divide the effective total volume of filaments by the cytoplasmic volume, assumed to be ~39 pL. This gives us effective volume fractions of 25% for F-actin and 94% for VIFs.

| | F-actin | Vimentin |
|---|---|---|
| $M_w$ (kDa) | 42 | 54 |
| $c$ (mg/mL) | 3 | 3 |
| $N$ (nm$^{-1}$) | 0.74 | 0.37 |
| $L_p$ (nm) | 17000 | 1000 |

**Fluorescence images of cell lines with vimentin labeled green**

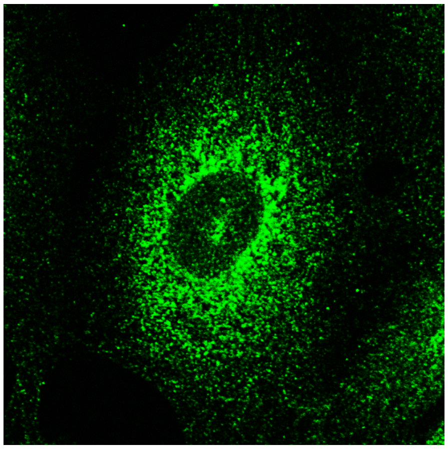

MEFs with a Y117L mutation, which allows lateral assembly of monomers into unit length filaments (ULF) but prevents the end-to-end annealing that forms long VIF

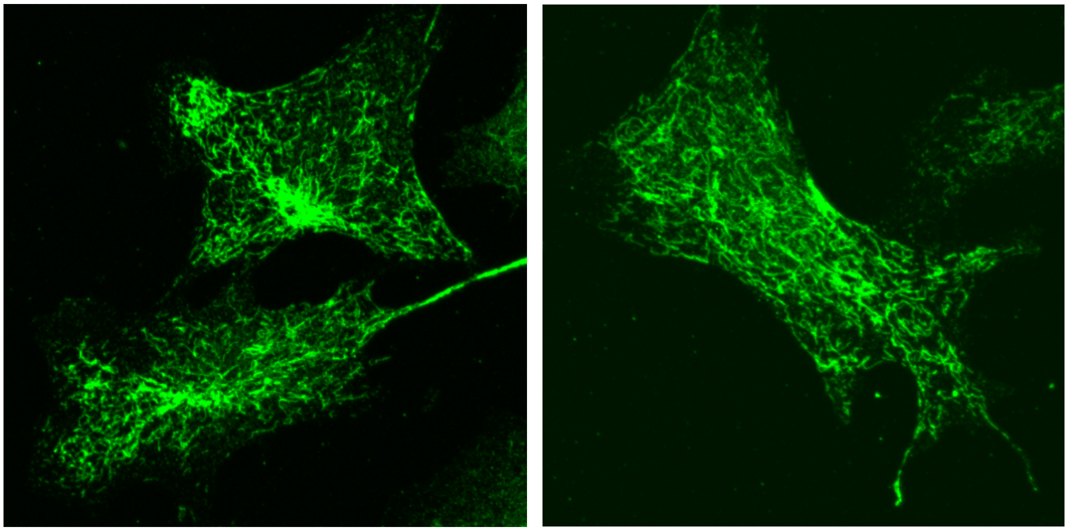

Vim-/- MEFs rescued by transfection with vimentin cDNA, which assemble long, mature VIFs, but whose vimentin protein expression is only ~30% of that of the endogenous concentration in WT MEFs
